# Functional corticostriatal connection topographies predict goal directed behaviour in humans

**DOI:** 10.1101/169151

**Authors:** Andre F. Marquand, Koen V. Haak, Christian F. Beckmann

## Abstract

Anatomical tracing studies in non-human primates have suggested that corticostriatal connectivity is topographically organized: nearby locations in striatum are connected with nearby locations in cortex. The topographic organization of corticostriatal connectivity is thought to underpin many goal-directed behaviours, but these topographies have not been completely characterised in humans and their relationship to uniquely human behaviours remains to be fully determined. Instead, the dominant approach employs parcellations that cannot model the continuous nature of the topography, nor accommodate overlapping cortical projections in the striatum. Here, we employ a different approach to studying human corticostriatal circuitry: we estimate smoothly-varying and spatially overlapping ‘connection topographies’ from resting state fMRI. These correspond exceptionally well with and extend the topographies predicted from primate tracing studies. We show that striatal topography is preserved in regions not previously known to have topographic connections with the striatum and that many goal-directed behaviours can be mapped precisely onto individual variations in the spatial layout of striatal connectivity.

---

A large body of work in experimental animals has shown that corticostriatal connectivity mediates a wide repertoire of goal-directed behaviors.^1-10^ This repertoire extends beyond the classical role of the striatum in planning and executing motor behaviour ^4^ and includes many other domains including reward processing,^5,6^ executive function,^7^ emotion^8^ and decision making.^7,9^ This has contributed to an emerging view of the striatum as a crucial locus for functional integration ^10,11^ and as a key region implicated in many brain disorders including Parkinson’s disease, obsessive compulsive disorder, attention-deficit hyperactivity disorder and schizophrenia.^12–14^

The striatum has extensive connections with virtually the entire cerebral cortex^15^ and an early influential theory of corticostriatal circuitry suggested that the multiple behaviours it subserves can cleanly be segregated into parallel segregated circuits.^3^ However, anatomical tracing studies in non-human primates have suggested that this view is too simplistic.^1,2,4,10,16^ These studies involve injecting anatomical tracers into predefined regions in the striatum or prefrontal cortex and mapping their terminal projection fields. On the basis of these results, it is thought that the projections from cortex are topographically organized in a ventral-to-dorsal and medial-to-lateral gradient across the striatum^1,2,10,16^ such that neighbouring locations in the striatum are connected to neighbouring locations in cortex.^1,2,4^ This topography does not appear to respect anatomical boundaries; for example, there is no clear boundary between ventral and dorsal striatum^11^ and the distinction between the caudate and putamen is primarily anatomical, whereas the distribution of cortical projections varies smoothly across both structures.^11,17^ Moreover, the projection zones from different cortical regions overlap within the striatum^10,16,18,19^ such that only cortical areas separated by large distances have completely non-overlapping striatal projection zones.^20^ The anatomical tracing approach has been invaluable in helping to understand corticostriatal circuitry because it can precisely localize the terminal fields of a given injection site. However, it cannot be applied to humans due to its invasive nature and it does not provide a quantitative map of the underlying connection topography; instead this must be inferred post-hoc from the distribution of terminal fields and – most importantly – it is fundamentally an anatomical measure and does not directly inform about function. This is particularly important in view of the broad relevance of corticostriatal circuitry to uniquely human behaviours and to brain function in health and disease.

Topographic representations are hallmark features of brain organization^21^ and while they have been well-studied in some brain systems (e.g. vision), their function outside sensory regions is poorly understood^21^ and few methods have been proposed to study them directly.^22^ Instead, the predominant approach has been to estimate connectivity between hard parcellations that define regions of interest in the striatum, cortex, or both. Typically this is done either on the basis of diffusion tensor imaging (DTI),^16^’^18,23-25^ resting state fMRI^26,27^ or meta-analytic data.^28^ While this approach has provided qualitative evidence for homology with the topography predicted by studies in experimental animals, it suffers from limitations: first, it does not provide quantitative measures of the overall topological organization that can be related to behaviour in individual subjects.

Second, the parcellation approach assumes that activity is constant across relatively large brain areas and is not well suited to studying topographic representations, where the connectivity varies gradually across space. Third, parcellations cannot – by definition – accommodate overlapping representations within the striatum. These problems are particularly acute in the striatum in view of the gradual connectivity pattern suggested by tracing studies, the convergence of projections from widespread cortical regions and the sheer breadth of behaviours that this circuitry underpins.

In this work, we aimed to estimate quantitative representations of the functional topography of corticostriatal circuitry in humans and to determine the behaviours those representations map onto. First, we capitalize on a recent methodological development^29,30^ to accurately estimate smoothly-varying and overlapping topographic connection patterns (‘connection topographies’) in the striatum on the basis of connectivity of each striatal location with the rest of the brain. We estimate these connection topographies quantitatively in individual subjects from resting fMRI. We then re-map these topographies across the cortex to provide detailed topographic maps of human corticostriatal circuitry. Finally, we chart the behavioural determinants of individual variations in these topographic representations across a battery that spans behaviours that depend on striatal function, including reward, executive function and psychopathology. We aimed to address four key questions: (i) can connection topographies in the striatum be quantified in humans at the level of individual subjects? (ii) How does the connection topography in the striatum and its remapping across the cortex differ from the topography predicted from experimental animals? For example, there is evidence from non-human primates that the temporal and parietal cortices project topographically to the striatum,^4,15^ but tractography studies in humans have not provided evidence for these projections;^18,23,25^ (iii) which behaviours do individual variations in these connection topographies map onto? and finally, (iv) do they map behaviour more reliably than parcellation-based approaches?

First, we employed resting-state fMRI from 466 subjects from the Human Connectome Project (HCP)^31^ to reconstruct connection topographies that map the connectivity of each striatal location with the rest of the brain. For this, we used an analysis approach^29,30^ that provides reproducible and parsimonious representations of overlapping connection topographies at the level of individual subjects (see Methods and Figure 4). Here, we restrict our analysis to two overlapping topographies but the approach can in principle be extended to capture further overlapping representations. This analysis approach is summarised in Figure 4 and described in the methods. Briefly however, it involves three main steps: first, we derive a similarity matrix that describes the similarity of the connection pattern (‘fingerprint’) of each striatal voxel with the rest of the brain. For this we choose the η^2^ coefficient.^29^ This step involves a lossless dimensionality reduction using singular value decomposition (SVD). Second, we feed this matrix into a manifold learning algorithm (Laplacian eigenmaps ^32^) to derive a set of connection topographies. Third, we fit a spatial statistical model (a ‘trend surface model’^33^) to each topography. This yields a set of coefficients for each subject providing a low dimensional representation of the connection topography that can be used for statistical analysis. Finally, we re-map connection topographies to the cortex to determine how the topographic connectivity pattern within the striatum is preserved across connections with the cortex. In our data, the trend surface models summarising the topographies were very accurate, explaining a mean (std. dev.) of 94.4% (2.2%) of the variance in the connection topographies across subjects and scanning sessions. These coefficients were also highly reproducible; we estimated separate models for each of the two repeated fMRI runs for each subject, which were highly correlated across runs (Left: mean (std. dev) ρ = 0.98 (0.05); Right ρ = 0.98 (0.04)). We illustrate both the variations across subjects and the reproducibility within subjects in Supplementary Figure 1.

The dominant connection topography corresponded strongly to the underlying anatomy both in a continuous sense (Supplementary Figure 2a) and after applying K-means clustering (Supplementary Figure 2b) such that the resulting group-level parcellation recapitulated the anatomical boundaries of the caudate, putamen and nucleus accumbens (Supplementary Figure 4). In contrast, the second overlapping topography exhibited a more gradual pattern of connectivity change corresponding with the dorsal-to-ventral, medial-to-lateral and anterior-to-posterior anatomical gradient of connectivity within the striatum predicted from tracing studies in non-human primates ^1,2^ (Supplementary Figure 2c). For both topographies, this correspondence was also evident in the individualized topographies (Supplementary Figure 1). Because of the prominence of this gradual connectivity pattern in the animal literature, and the overwhelming focus on parcellation in the human literature, we focus the remaining analysis on the second topography.

Next, we examined the connectivity profile of the second topography by remapping this representation from the striatum onto the cortex. We achieved this by color-coding each cortical vertex according to the striatal voxel that it correlates the most with.^22,29^ In experimental animals, a gradient of topographical connectivity in the striatum has been most frequently associated with reward, where the projections from many reward-related brain regions converge in the rostral striatum.^5^ Therefore, we examined the correspondence between the topographic representation in the striatum and these reward related areas. The connectivity profile between the ventral rostral striatum and medial prefrontal cortex and midbrain showed a striking similarity to the connection topography predicted from invasive tracing studies (Figure 1a).

**Figure 1:**
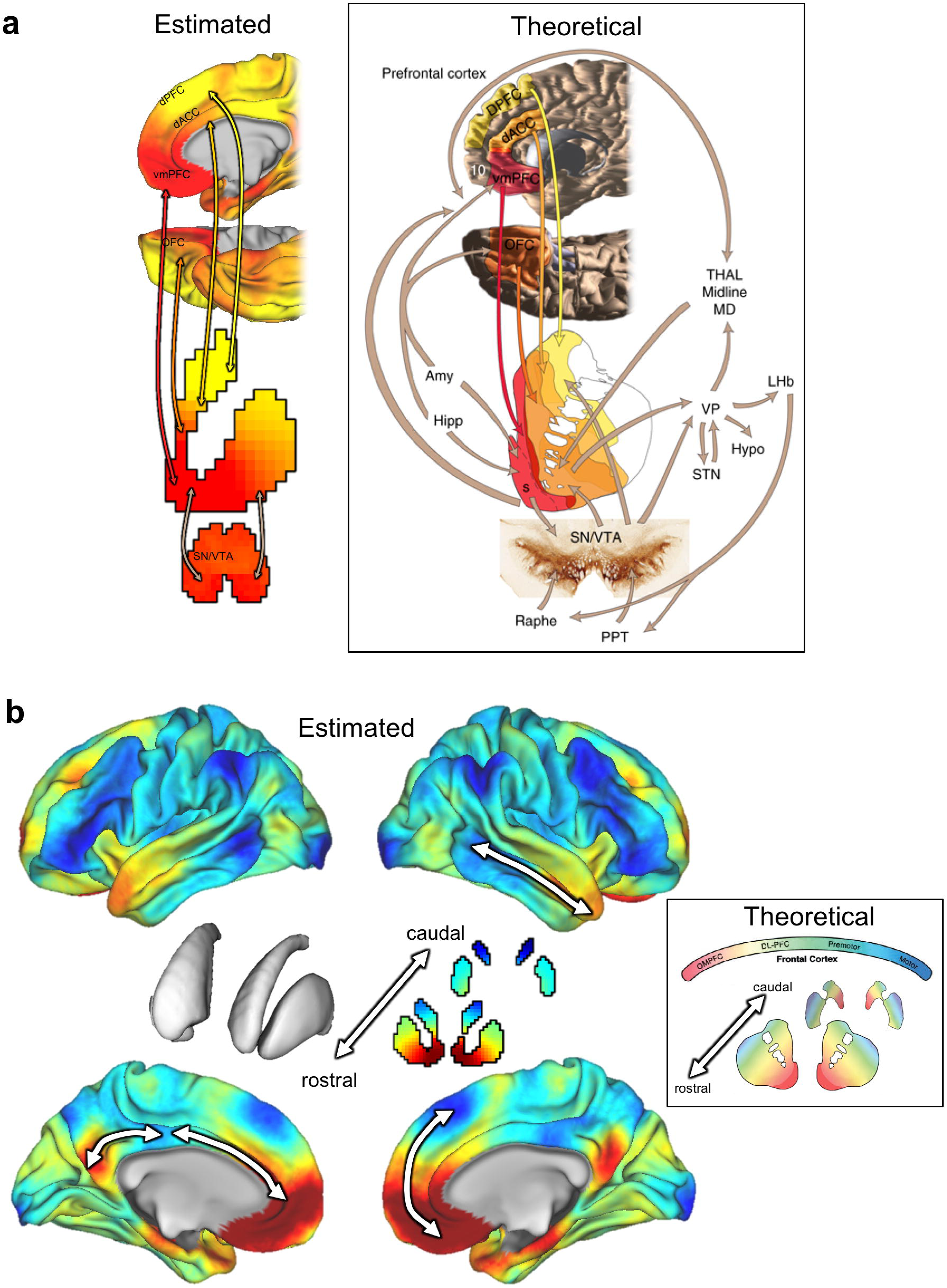
Panel a: The striatal connection topography estimated here from resting state fMRI in humans (Left)shows an excellent correspondence with the theoretical pattern of connectivity between reward-related brain areas and the striatum based on invasive tracing studies in animals (Right, reproduced from ^5^). Panel b: The pattern of connectivity between the connection topography in the striatum (centre) and the cerebral cortex. Arrows indicate examples of cortical regions for which topographic connectivity with the striatum is preserved. The inset shows the theoretical connectivity pattern derived from invasive tracing studies in non-human primates (modified from ^1^). Note that the colour scheme in panel b has been changed relative to panel a to match the existing non-human primate literature^1,2,11^ Abbreviations: dPFC: dorsal prefrontal cortex; DL-PFC = dorsolateral prefrontal cortex; vmPFC = ventromedial prefrontal cortex; OFC = orbitofrontal cortex; SN/VTA = substantia nigra/ventral tegmental area; Hipp = hippocampus; Amy = amygdale; STN = subthalamic nucleus; OMPFC = orbitomedial prefrontal cortex; Raphe = Raphe nucleus; PPT = penduclo-pontine tegmentum; Flypo = hypothalamus; Thal Midline MD = mediodorsal thalamus; VP = ventral pallidum; LHb = lateral habenula.

Reward is a primary behaviour dependent on corticostriatal circuitry, through its connections with cortical areas involved in reward processing. However, the striatum has extensive connections with nearly all cortical areas. Therefore, we next investigated the broader topographic connectivity profile of the striatum. Again, the re-mapped topographic connectivity pattern shows an excellent correspondence with the pattern of projection targets that has been predicted from tracing studies in non-human primates (Figure 1b). Whilst this correspondence is reassuring, we emphasize that these re-mapped topographic representations were obtained from humans in vivo, extend across all cortical areas and in considerably more detail than is provided by the theoretical model. Most importantly, they are also quantitative and can therefore be related to human behaviours in a statistical manner.

These results show that human corticostriatal connection topographies extend beyond the known pattern of topographical connectivity derived from studies of non-human primates and span multiple spatial scales. Studies of corticostriatal circuitry in experimental animals have mostly focused on prefrontal cortex^11^ although some studies have provided evidence that the striatum is topographically connected with the parietal and temporal lobes.^15^ The remapping of the striatal connection topographies across the cortex shows that the topography in the striatum was recapitulated across many cortical regions (Figure 1b, arrows), which suggests that many cortical regions have topographically organized connections with the striatum. Some of these connections are expected from animal studies, for example: the amygdala, hippocampus and ventromedial prefrontal cortex showed a strong correspondence to ventromedial striatal regions, which in monkeys is the only area of the striatum that receives input from these structures^11^ Similarly, cortical regions associated with motor function (e.g. primary- and supplementary motor areas) showed the expected correspondence to the lateral putamen. On the other hand, other features have not been described in the animal literature. For example, the posterior cingulate cortex was strongly connected with the ventromedial striatum and lateral prefrontal cortex showed a strong correspondence to dorsal and posterior caudate. Also in contrast to the non-human primate literature, we also found that the gradient of connectivity within the striatum could be traced onto rostral-to-caudal gradients within connected brain regions. For example, we detected connectivity gradients within the anterior cingulate, posterior cingulate and temporal cortices (Figure 1b).

Finally, we were interested in determining the correlates of these connection topographies across a wide range of goal-directed behaviours.^11^ Therefore, we employed a unified multivariate analysis approach to find associations between the full set of trend surface model coefficients from each hemisphere and the extensive battery of behavioural measures derived from the HCP dataset. This battery includes measures of many aspects of cognition, reward, language, emotion, personality and clinical scales across multiple diagnostic categories.^34^ Specifically, we used canonical correlation analysis (CCA), a multivariate analysis technique that seeks patterns of covariation between datasets (see Methods). In both hemispheres, we detected a highly significant association between interindividual variations in the second topography and the behavioural battery (left hemisphere: ρ = 0.74, p < 0.002 (permutation test), Wilk’s Lambda = 0.0094, p < 0.005; right hemisphere: ρ = 0.73, p < 0.01, Wilk’s Lambda = 0.009, p < 0.002). Since we considered it to be unlikely that this association could be cleanly partitioned into orthogonal components (an inherent feature of the CCA decomposition), we mapped the behavioural domains underlying this association across the entire decomposition (Figures 2 and 3). These figures show structure coefficients corresponding to all of the 9 canonical components from the CCA decomposition (9 because the optimal statistical model was a polynomial of model order 3). These provide a measure of importance of each brain voxel (Figures 2a and 3a) or behavioural score (Figures 2c and 3c) in maximizing the correlation between brain topography and behaviour. Successive components show additional contributions to maximizing the correlation orthogonal to the other components (see Methods for details). This showed that the association in the left hemisphere was driven principally by: (i) delay discounting, consistent with the known role of the striatum in reward and delay valuation^7^ (ii) relational processing^34^ which is relevant because the task they were derived from was designed to precisely localize rostrolateral prefrontal cortex^34,35^ and (iii) psychological wellbeing, which is also plausibly related to corticostriatal function.^34^ In the right hemisphere, the association was driven by: (i) social cognition, derived from a task that elicits robust activations in brain areas associated with theory of mind^34^ (ii) sustained attention, which is known to depend on corticostriatal circuitry^7^ and is consistent with its role in the pathophysiology of attention deficit/hyperactivity disorder^12^ and (iii) personality, which also depends on corticostriatal connectivity.^24^ To assess reproducibility, we repeated the entire pipeline on the second fMRI run. The pattern of results was largely consistent with that derived from the first fMRI run: associations were detected for both hemispheres (left: ρ = 0.74, p < 0.001, Wilk’s Lambda = 0.013 (not significant); right: ρ = 0.75 p < 0.001, Wilk’s Lambda = 0.009, p < 0.004). Note that although the entire decomposition did not reach significance for the left hemisphere, individual predictive gradients did. The particular behavioural domains underlying these associations were also similar (Supplementary Figures 4 and 5), where again delay discounting and relational processing were strongly associated with the connection topography in the left hemisphere, and sustained attention was dominant in the right hemisphere. However, there were also some differences in the behavioural measures associated with the connection topographies; for example, emotional processing scores from the fMRI task were associated with the connectivity gradients in the second fMRI run but not the first (see below).

**Figure 2:**
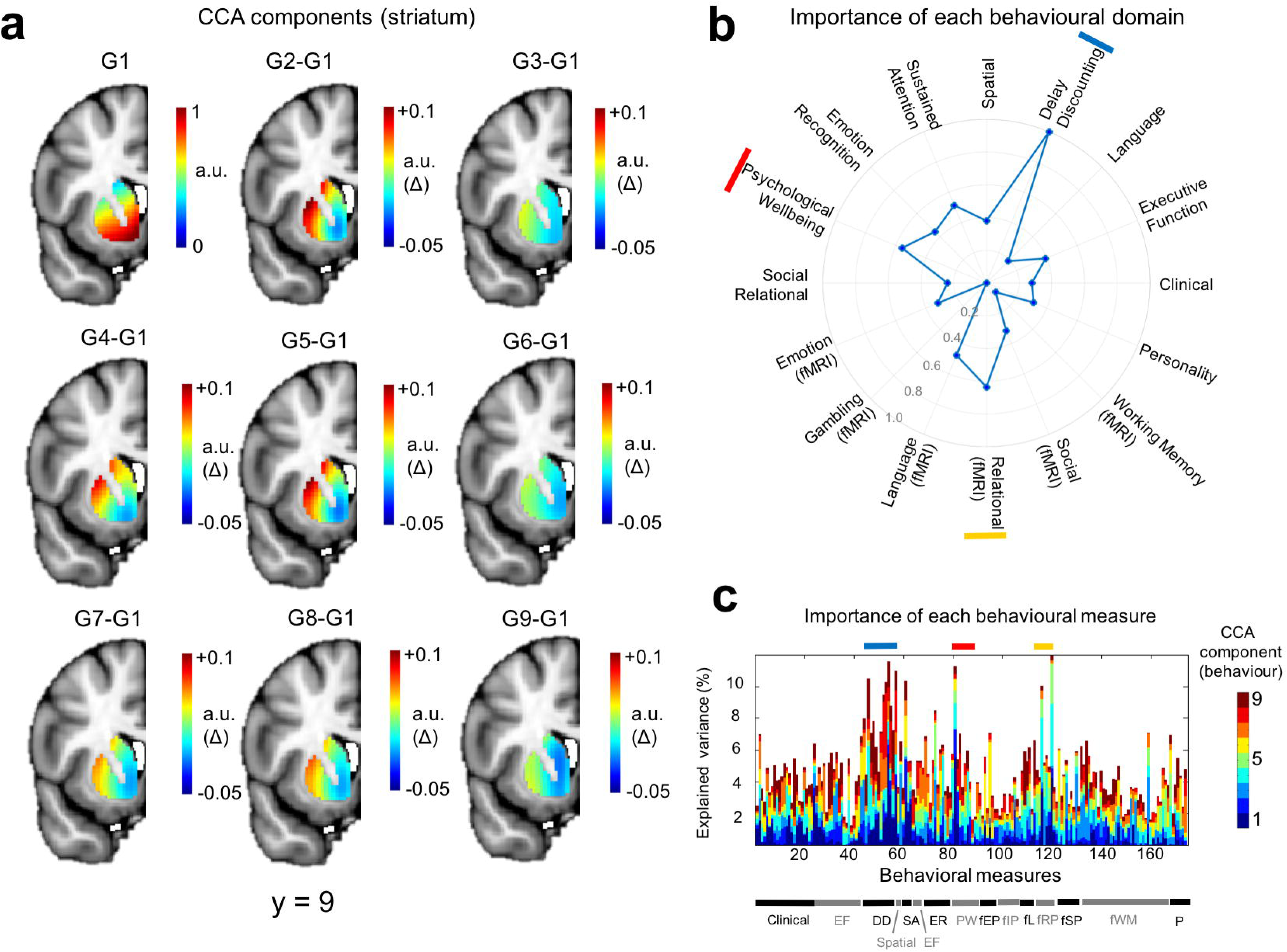
Relative importance of different variables in driving the multivariate correspondence between the topography from the left striatum and the behavioural battery (derived from all 466 subjects used in the analysis). Panel a: Importance of connectivity gradients in predicting the behavioural scores. The top left panel (G1) shows structure coefficients corresponding to the principal predictive gradient estimated by CCA. These are rescaled such that the maximum in the image is equal to one and can therefore be considered to be in arbitrary units (a.u.). The remaining panels show differences between the rescaled principal CCA gradient and each successive rescaled predictive gradient (a.u. delta). For example, G2-G1 is the difference between the second gradient and the first. This helps to highlight the differences between the predictive gradients. Panel b: Predictive pattern of measures contributing to the CCA predictions. These are the structure coefficients aggregated across behavioural domains (see Methods) and are also rescaled such that the maximum behavioural domain has a value of 1, here represented by a point on the outermost circle. The top three domains are indicated by red, yellow and blue bars. Panel c: Structure coefficients for all of the 174 individual behavioural items. Behavioural domains are indicated by the bar at the bottom and a full list of individual items is provided in the supplementary Methods. Coloured bars indicate the top three domains (see panel b). Abbreviations: CCA = canonical correlation analysis; DD = Delay discounting; EF = executive function; fEP = emotion processing (fMRI); fIP = incentive processing (fMRI); fL = Language (fMRI) fRP = relational processing (fMRI); fSP = social processing; fWM = working memory (fMRI); ER = emotion regulation; PW= psychological wellbeing; SA = sustained attention; P = personality.

**Figure 3:**
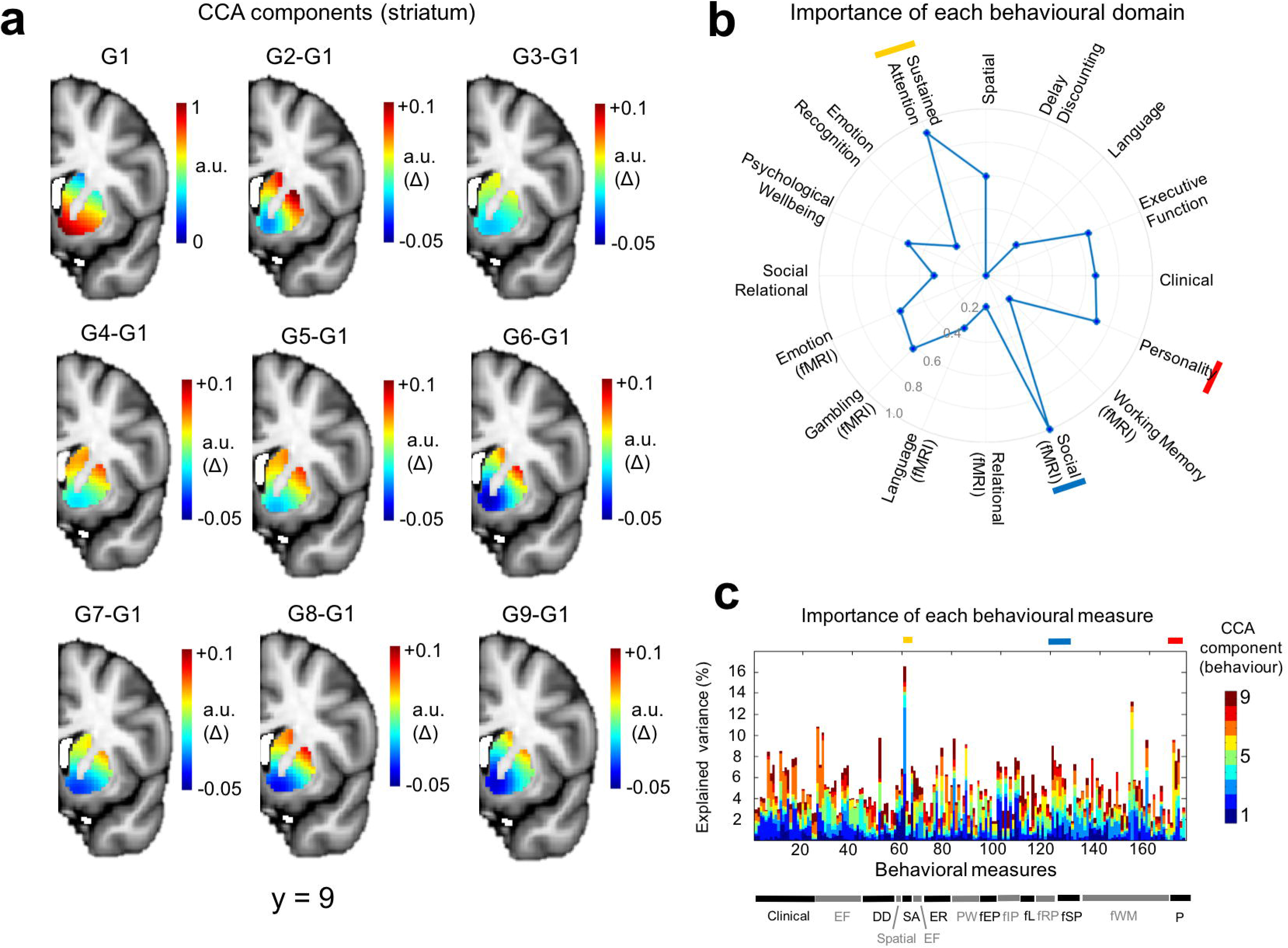
Relative importance of different variables in driving the multivariate correspondence between the topography from the right striatum and the behavioural battery (derived from all 466 subjects used in the analysis). Panel a: Importance of connectivity gradients in predicting the behavioural scores. The top left panel (G1) shows structure coefficients corresponding to the principal predictive gradient estimated by CCA. These are rescaled such that the maximum in the image is equal to one and can therefore be considered to be in arbitrary units (a.u.). The remaining panels show differences between the rescaled principal CCA gradient and each successive rescaled predictive gradient (a.u. delta). For example, G2-G1 is the difference between the second gradient and the first. This helps to highlight the differences between the predictive gradients. Panel b: Predictive pattern of measures contributing to the CCA predictions. These are the structure coefficients aggregated across behavioural domains (see Methods) and are also rescaled such that the maximum behavioural domain has a value of 1, here represented by a point on the outermost circle. The top three domains are indicated by red, yellow and blue bars. Panel c: Structure coefficients for all of the 174 individual behavioural items. Behavioural domains are indicated by the bar at the bottom and a full list of individual items is provided in the supplementary Methods. Coloured bars indicate the top three domains (see panel b). Abbreviations: CCA = canonical correlation analysis; DD = Delay discounting; EF = executive function; fEP = emotion processing (fMRI); fIP = incentive processing (fMRI); fL = Language (fMRI) fRP = relational processing (fMRI); fSP = social processing; fWM = working memory (fMRI); ER = emotion regulation; PW= psychological wellbeing; SA = sustained attention; P = personality.

To determine whether these behavioural associations were better captured by a gradual connection topography relative to piece-wise constant parcels, we performed additional analyses, where we repeated the CCA analysis after applying K-means to generate an individualized parcellation for each subject (i.e. similarly to the group-level parcellation in Figures S3a and S4) then averaging the principal gradient across each parcel. In both cases, we detected some associations between behaviour and connectivity, but these were slightly less reproducible across fMRI runs. More specifically, significant associations were detected for both left and right hemisphere, but each in only one of the fMRI runs (first run, left hemisphere: ρ = 0.64, Wilk’s Lambda = 0.27, both not significant; first run, right hemisphere: ρ = 0.80, Wilk’s Lambda = 0.14, both p < 0.001; second run, left hemisphere: ρ = 0.70, p < 0.007, Wilk’s Lambda = 0.21, p < 0.002; second run, right hemisphere: ρ = 0.67, Wilk’s Lambda = 0.21, both not significant). Note that this approach provides an optimistic estimate of the association achievable through parcellation because the parcellation is applied after employing manifold learning which provides the advantage of separating the signal attributable to overlapping representations and is not typically done for parcellation approaches.

To summarise these results: we estimated smoothly varying patterns of corticostriatal connectivity non-invasively from humans, which: (i) show that connection topographies can be reliably identified at the level of individual subjects; (ii) provide quantitative topographic maps of human corticostriatal circuitry that show an excellent correspondence to the topography predicted from animal tracing studies^1,2,5,11^ whilst also showing that striatal topography is preserved in brain regions that have not been shown to have topographic connections with the striatum either in humans (e.g. temporal and parietal lobes) or experimental animals (e.g. posterior cingulate cortex). They also allowed us to demonstrate that: (iii) individual variations in these topographic connectivity patterns predict specific behaviours in a way that is (iv) more reproducible than a parcellation-based approach.

The unique feature of our approach is that we shift away from hard parcellation and instead perform inferences directly on the basis of spatial connection topographies. This is particularly well-suited to studying corticostriatal function because it provides inferences at the level of overlapping and spatially distributed connectivity patterns across the striatum, not at the level of piece-wise constant parcels. Our results challenge the dominance of the parcellation view of brain connectivity, and our approach overcomes several methodological problems it entails: our topographic approach accommodates multiple connectivity patterns that overlap in the same structure, it does not require defining the number of parcels or the parcellation strategy in advance, which is important because the parcel is the fundamental unit of most network-based approaches to connectivity, and therefore any errors in parcel definition are propagated through the entire analysis. This provides important advantages over existing methodology because: (i) the number of parcels is often not well-defined^28^ and (ii) purely anatomical parcellations often do not map well onto function^36^ whereas (iii) there are wide range of data-driven parcellation strategies, which often show only a moderate correspondence with one another.^27,28^ These problems notwithstanding, our approach is complementary to parcellation strategies; in the striatum, parcellation is a useful approach for detecting segregated parallel circuits^18^ or identifying their projection zones^16^ but do not lend themselves naturally to making inferences about convergent processing because they do not easily accommodate the overlapping representations that are fundamental to corticostriatal circuitry.^1,5^ In our data, we were also able to detect behavioural associations with an individualized parcellation of the striatum, but these were less reproducible than the continuous connection topographies. This suggests that the parcellation may coarsely approximate the underlying continuous topography.

Similar to other studies that have employed functional connectivity to investigate corticostriatal circuitry,^26,27^ our results should be interpreted in the context of the use of a functional connectivity method that is sensitive to both direct (i.e. monosynaptic) and indirect (polysynaptic) connections. This provides advantages and disadvantages relative to the predominant approach to studying structural corticostriatal connectivity in humans on the basis of DTI.^16,18,23-25^ The advantage of structural connectivity methods is that they provide a means to directly identify monosynaptic connections between brain regions, but there is no guarantee that the presence or absence of a given structural connection is functionally relevant. It is also well-known that DTI-based methods require myelinated connections and can have difficulties in following fibres through areas with extensively crossing fibre bundles. On the other hand, an advantage of the present approach is that it provides a means to identify functionally connected brain regions even if they are not connected monosynaptically or if tracts between them cannot be identified. In some settings, functional connectivity can be used to identify monosynaptic connections on the basis of partial correlations between brain regions. However, this requires that the relevant brain area first be subdivided into atomic units (e.g. via parcellation). This is a reasonable approach in many cases, but partial correlation is not directly applicable here because we assess the similarity of connectivity patterns at the level of individual voxels and data within the striatum and within cortical areas are inherently smooth such that neighbouring voxels are highly correlated.

Our results that validate and extend the literature based on experimental animals in three ways: first, ventromedial striatal areas were strongly connected to networks that subserve reward (e.g. ventromedial and orbitofrontal cortex, dorsal anterior cingulate, amygdala and hippocampus). This is in line with studies that have shown that projections from reward-related brain regions overlap most extensively in the rostral striatum^5^ and with studies in experimental animals that suggest that the amygdala and hippocampus are important nodes in the extended reward network.^5,37^ Behaviourally, this was reflected in an association between the connectivity patterns and delay discounting, which extends studies that have principally demonstrated a group-level association between delay discounting and mean activity in the ventral striatum.^38^ Second, dorsal and caudal striatal regions were strongly connected to lateral prefrontal regions, which is in line with evidence from nonhuman primates showing that terminals from the motor and premotor striatum do not extend into rostral striatum.^11^ In humans, this is also consistent with the involvement of the dorsal striatum in executive function and decision making^7^ and with observations that lesions in the dorsal caudate nucleus cause impairments in working memory.^39^ Third, lateral striatal regions (e.g. mid putamen) were strongly connected with motor regions, as would be expected based on the well-documented role of the putamen in motor circuitry.^4^

Our data also provide evidence for the existence of topographic connections that would not have been predicted on the basis of the existing literature, which has predominantly focused on projections that pass through prefrontal cortex.^3^ For example, the posterior cingulate cortex was topographically connected with ventral striatum. The posterior cingulate cortex has a well-documented role in brain’s default mode network (DMN) which shows reduced activity during many goal-directed behaviors.^40^ This is of particular relevance to the associations we detected with delay discounting and sustained attention because there is increasing evidence that the striatal dopamine system modulates the DMN^37,41^ and failures in suppression of DMN activity have been associated with momentary lapses in attention.^42^

Another key finding was evidence for a topographic gradient of connectivity within the temporal and parietal lobes. This is consistent with evidence of topographically organized patterns of connectivity between striatum and temporal and parietal cortices in experimental animals.^19,43,44^ However, these gradients have not, to date, been reported in studies that have studied human corticostriatal circuitry.^18,23,25^ This may be because these human studies all used structural connectivity methods (e.g. tractography). This is particularly relevant because fibres from parietal and temporal cortices need to pass through areas of complex fibre crossings to reach the striatum.^25^ In our data, connectivity with the temporal lobe may be related to some of the other behaviours associated with the striatal connection topographies (e.g. social functioning).

Our results are broadly consistent with other studies which have employed functional connectivity approaches to identify the connectivity of the striatum.^26,27^ Di Martino and colleagues used seed-based connectivity to identify cortical voxels correlated with each of a set of anatomically defined regions of interest.^26^ Choi and colleagues applied a parcellation approach to divide the striatum into subregions by assigning each striatal voxel to its most strongly correlated cortical network.^27^ These studies provided evidence for the existence of a seed-location dependent functional difference in striatal organization. Our approach is complementary to these studies and enabled us to: (i) quantify the functional topography in the striatum directly and in a smoothly varying manner; (ii) provide a low-dimensional representation of this topography that can be related quantitatively to behaviour and (iii) disentangle overlapping representations within the striatum. Here, this was reflected as a gradient of functional connectivity superimposed on the underlying anatomy (Supplementary Figure 2). This latter property is particularly important in the striatum given the sheer number of behaviours that corticostriatal circuitry underpins in humans.

While our results suggest that the association with delay discounting was lateralized in that the association was most prominent in the left hemisphere, we are cautious of such an interpretation because there was weak evidence of an association between delay discounting and the dominant connection topography after parcellation that did not achieve statistical significance. An avenue for further study is to investigate the stability of the behavioural associations over repeated measurements. Although the most important behavioural associations were reproducible across runs (e.g. the association with delay discounting), there was also some run-to-run variability in the particular behavioural variables underlying the brain-behaviour correspondence. The most salient of these was that measures derived from the fMRI emotional processing task were associated with the connection topographies in both hemispheres from the second resting fMRI run but not the first.

This may be because the emotional task was acquired in the same scanning session as the second fMRI run. According to the HCP protocol (https://www.humanconnectome.org/documentation/data-release/Ql_Release_Appendix_1.pdf). three of the fMRI tasks were acquired after the first resting fMRI run and four were acquired after the second run (including the emotion processing task). Emotion processing is dependent on corticostriatal circuitry^8^ and emotional task performance is correlated with dopamine release in the striatum.^45^ Therefore, we speculate that state-dependent effects related to corticostriatal circuitry were related to emotion processing. If correct, this hypothesis underscores the importance of considering multiple measurement timepoints to detect state-dependent effects. In future work we will also investigate methods to infer directionality of the connection topographies, which remains a challenging problem for most approaches to estimation of functional connectivity.^46^ Finally, another open question is determining the model order for overlapping connection topographies. Here we considered the first two overlapping topographies, but it is also likely that higher order topographies could provide information at finer levels of detail.

In summary, we mapped topographic connectivity between striatum and cortex in humans. Our results simultaneously correspond with and extend the connection topographies predicted from studies in experimental animals. We demonstrate that topographic connectivity with the striatum in humans extends more widely than previously thought and we precisely map the behaviours that these topographies predict across an extensive behavioural battery. Our results lend support to the notion that the striatum functions as a hub for integrating information from cortical networks subserving many human goal directed behaviours and our approach provides a means to estimate continuous connection topographies at the level of individual subjects and to relate individual variations in these topographies quantitatively to behaviour.

## Methods

### Neuroimaging data

Resting state fMRI data were derived from the Human Connectome Project (HCP), which aims to acquire exceptionally high-quality neuroimaging data from more than 1000 twins and non-twin siblings.^31^ Full details surrounding the sample, data acquisition, ethical and preprocessing procedures have been reported previously,^31,34,47^ but in brief, our sample (N=466, aged 22-26, 293 females) included all subjects from the ‘500 subjects’ release who completed all resting fMRI scanning runs and for whom sufficient data were available to determine familial relationships. This was necessary to avoid introducing bias in the non-parametric statistical inference procedures we employ (below). We anticipated that this sample would be sufficient to identify salient multivariate brain-behaviour relationships based on previous results using a similar analysis approach.^48^ Participants were scanned on a customized 3 Tesla Siemens Skyra scanner (Siemens AG, Erlanger, Germany) using multi-band accelerated fMRI four times over two days, with each run comprising 15 min. Resting fMRI data were preprocessed according to the HCP minimal processing pipeline^47^ then denoised using advanced artefact removal procedures based on independent component analysis^49^ before being smoothed with an 6mm kernel that respected the geometry of the brain. Specifically, subcortical structures were treated volumetrically, while cortical structures were projected onto the cortical surface according to the documented HCP procedures.^47^

### Estimation of connection topographies

We estimated connection topographies from the HCP resting state fMRI data separately for each subject, hemisphere and fMRI scanning session. For this, we used an emerging approach that enables the dominant modes of functional connectivity change within the striatum to be traced on the basis of the connectivity between each striatal voxel and the rest of the brain. This procedure is described elsewhere^29,30^ and is summarized in Figure 4. Briefly, we rearranged the fMRI time series data from the both the striatum and all grey-matter voxels outside the striatum into two time-by-voxels matrices. Since the latter is relatively large, we losslessly reduced its dimensionality using singular value decomposition (SVD). We then computed the correlation between the voxel-wise striatal time series data and the SVD-transformed data from outside the striatum, then used the η^2^ coefficient to quantify the similarities among the voxel-wise fingerprints (see ^29^). Then, we applied the Laplacian eigenmaps manifold learning algorithm^32^ to the resulting similarity matrix, resulting in a series of vectors that represent the dominant modes of functional connectivity change (i.e. connection topographies). Note that this can be done at the group-level by using the average of the individual similarity matrices (as in Supplementary Figure 3) or individually for each subject (as used for statistical analysis). In the latter case, the resulting connection topographies were highly consistent across fMRI runs (Supplementary Figure 2), in line with what we have demonstrated previously for other brain regions.^29^ Finally, to enable statistical analysis over these connection topographies we fit a spatial statistical model that provides an accurate representation of the topography in a small number of coefficients. For this, we use a ‘trend surface modelling’ approach^33^ which involves fitting a set of polynomial basis functions defined by the coordinates of each striatal location to predict each individual subject’s connection topography. We fit these models using Bayesian linear regression,^50^ where we employed an empirical Bayes approach to set model hyperparameters. Full details are provided elsewhere^50^ but this essentially consists of finding the model hyperparameters (controlling the noise- and the data variance) by maximizing the model evidence or marginal likelihood. This was achieved using conjugate gradient optimization. For fixed hyperparameters, the posterior distribution over the trend coefficients can be computed in closed form which, in turn, enables predictions for unseen data points to be computed. We used the maximum a posteriori estimate of the weight distribution as an indication of the importance of each trend coefficient in further analyses. To select the degree of the interpolating polynomial basis set, we fit these models across polynomials of degree 2-5 then compared the different model orders using the Bayesian information criterion. This criterion strongly favoured a polynomial of degree 3, which was taken forward for further analysis. Note that this decision was not strongly dependent on the choice of criterion because the Akaike information criterion resulted in identical conclusions.

**Figure 4:**
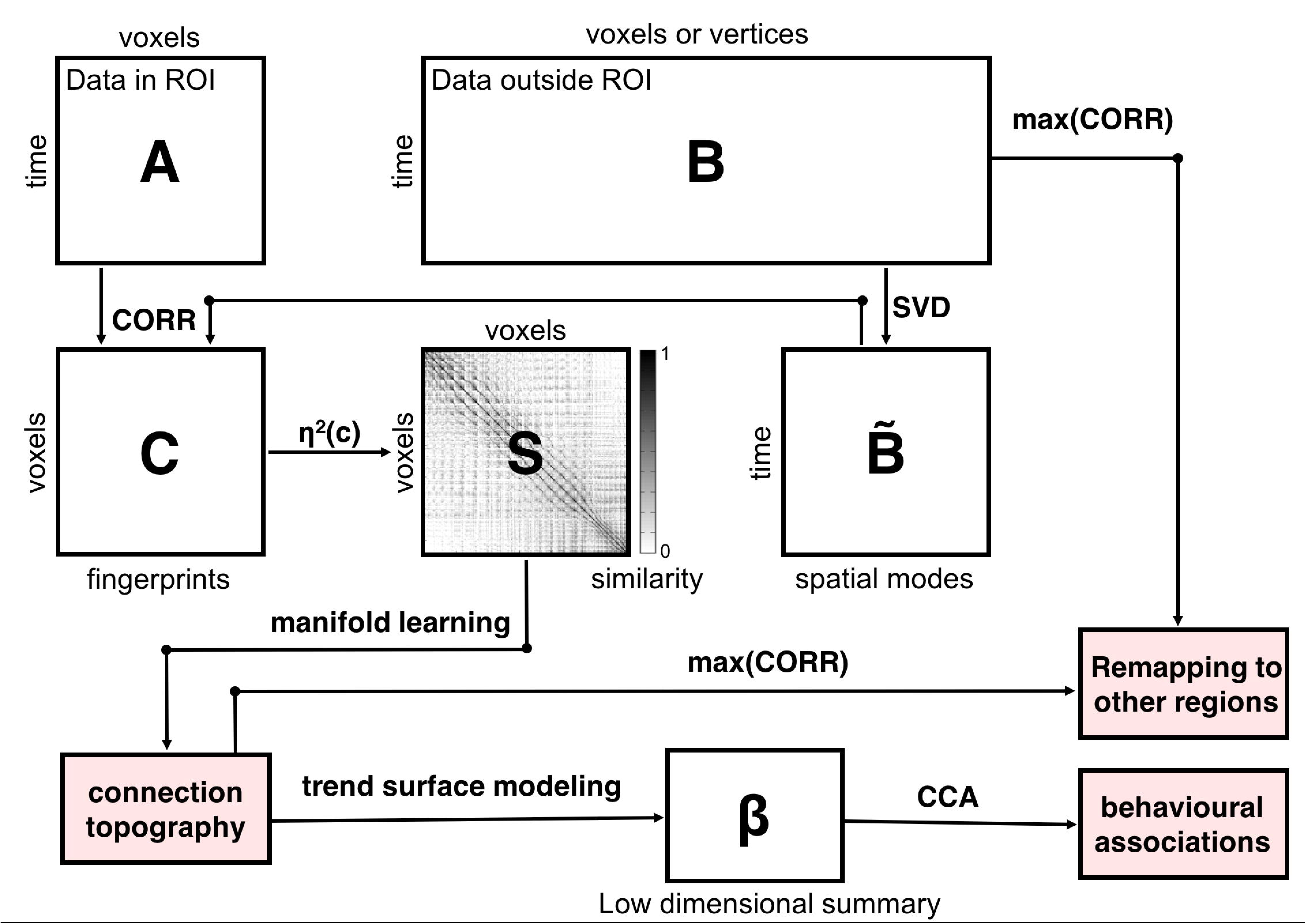
Summary of the analysis pipeline. The fMRI time-series data from a pre-defined region-of-interest (ROI) are rearranged into a time-by-voxels matrix A, as are the time-series from all vertices/voxels outside the ROI (matrix B). For reasons of computational tractability, the dimensionality of B is losslessly reduced using singular value decomposition (SVD), yielding B͂. For every voxel within the ROI, its connectivity fingerprint is computed as the Pearson correlation between the voxel-wise time-series and the SVD-transformed data, yielding matrix C. Then similarity between voxels is computed using the η^2^ coefficient. Manifold learning using Laplacian eigenmaps is then applied to this matrix, yielding a set of connection topographies, which can remapped to other regions by taking the maximum correlation. Then, trend surface modelling is applied to summarize these connection topographies by fitting a set of trend coefficients ((β) that optimally combine a set of spatial polynomial basis functions. Finally, canonical correlation analysis (CCA) is used to find associations with these behavioural measures. See Methods for further details

To determine the preservation of each striatal topography across its connections with other brain regions we performed a simple approach whereby we color-coded each cortical vertex or subcortical voxel according to the striatal voxel that it correlates the most with.^22^ This is ideal for our purposes because it is constrained to directly estimating corticostriatal interactions.

### Behavioural data

We evaluate the behavioural correlates of the connection topographies across an extensive battery of behavioural measures. This battery has been described elsewhere^31,34^ and includes demographic data (e.g. age, sex), psychometric data across multiple domains of functioning (e.g. cognition, emotion, personality, sensory processing and life functioning) plus clinical assessments spanning multiple psychiatric domains and diagnostic categories (e.g. substance use, impulse control, mood, anxiety and eating disorders).^34^ We also include behavioural measures from a set of fMRI tasks measuring working memory, incentive processing, motor function, language processing, relational processing, social cognition and emotion processing.^34^ In our analysis, we aimed to include the most extensive set of measures possible and therefore select variables using a similar strategy to a prior report using these data.^48^ Specifically, from the complete HCP battery^34^ (https://wiki.humanconnectome.org/display/PublicData/HCP+Data+Dictionary+Public+500+Subiect+Release) we included all psychometric data, clinical assessments and behavioural measures derived from the fMRI tasks but excluded basic demographic information, fields related to the data acquisition, and fields containing cortical thickness estimates derived from structural MRI. We also excluded physical motor variables (e.g. endurance and grip strength) and sensory processing variables (e.g. audition and olfaction), which we did not expect to be associated meaningfully with striatal function. We then removed data items with > 50% missing data or with a severe imbalance such that >95% of the subjects had the same value and used median imputation for the remaining missing data. Finally, we collected these variables into a matrix and ensured that this matrix was not rank deficient using Gaussian elimination. This procedure finds a basis for the range of the matrix by excluding a small number of variables (5-7 items in our data). This matrix was then used as input to the multivariate analysis described below. The final list of measures we employed is reproduced in the supplementary Methods. To assist interpretation, after analysis we grouped these into 14 categories, eight derived from established behavioural instruments (clinical, executive function, delay discounting, spatial orientation processing, sustained attention, emotion regulation, psychological wellbeing and personality), and six derived from behavioural measures recorded during the fMRI tasks (emotional processing, incentive processing, language, relational processing, social processing, working memory). See supplementary Methods for details.

To group the resulting scores for these variables (i.e. CCA structure coefficients – see below) into these categories, we used a simple Bayesian averaging approach that accommodates differences in the number of variables included in each category. This was desirable because the size of the categories was highly variable and some of the categories were very small. For a given random variable *Z*, this average is computed simply as
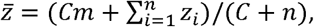
where *n* is the total number of data points, *m* is the prior mean and *C* is a constant equal to the average dataset size. Here, the mean was taken to be relatively non-informative and was set to 1/*n*. Although this yielded similar results to the ordinary arithmetic mean, it provides a more robust estimate of the contribution of the category as a whole because it reduces the chance that a category with a small number of variables scores highly because it contains (for example) a single informative variable.

### Statistical analysis

To test for an association between the behavioural battery and the connection topographies, we performed a single unified statistical analysis using canonical correlation analysis (CCA). CCA is a standard multivariate statistical approach that aims to learn a set of projection weights for each dataset that maximize the correlation between datasets. More concretely, given two data matrices, *X_n×p_* and *Y_n×q_* that have the same number of samples (*n*) but possibly different numbers of variables (*p* and *q*), CCA seeks canonical vectors *a* and *b* such that the projections *a’X* and *b’Y* are maximally correlated. These projections are referred to as the first pair of canonical variables. After this, CCA seeks each additional pair of canonical vectors which maximize the correlation between canonical variables, subject to the constraint that they are uncorrelated with the other pairs of canonical variables. Prior to analysis, we standardized the data in both datasets separately, which also ensured that the polynomial coefficients derived from the trend surface analysis were orthogonal. In contrast, the behavioural data exhibited strong multicollinearity which is known to cause problems with the interpretation of coefficients in linear models. Therefore, we inferred the association of each behavioural measure with the connection topography using structure coefficients. These are widely used in multivariate statistics to solve this problem^51^ and are defined as the univariate Pearson correlation between each measure and the predictions made by the CCA model. To infer the association of the connection topography with the behavioural scores, we compute forward maps^52^ for the canonical vectors. If the data are standardized as they are here, these are equivalent to structure coefficients.

To test the significance of the canonical correlation, we employed a permutation testing procedure that accounts for correlations induced by the familial relationships between subjects in the HCP sample.^53^ Specifically, we permuted the rows (i.e. subjects) of one of the data matrices 1000 times in a way that accommodates their familial relationships, computing the canonical correlation for each permutation. We then tested for significance by computing the centile of the non-permuted canonical correlation against an empirical null distribution derived from fitting a Gauss-Gamma mixture distribution to the permuted correlations^54^ This was necessary because in preliminary testing we found evidence that the reported familial structure did not fully account for the nuisance covariation structure between subjects. For each CCA decomposition, we used this procedure to test both the principal correlation coefficient and Wilk’s Lambda statistic. These statistics provide complementary information about the underlying CCA distribution: the principal correlation coefficient tests the magnitude of the dominant (or successive) mode(s) of canonical correlation, while Wilk’s Lambda tests the significance of the whole distribution.

### Data availability

All data used in this study are available to download from the Human Connectome Project website (http://www.humanconnectomeproiect.org/).

### Code availability

The computer code that support the findings of this study are available from the corresponding author upon reasonable request.

## Acknowledgments

We gratefully acknowledge support from the Netherlands Organization for Scientific Research (NWO) by VIDI grants to AFM (Grant No. 016.156.415) and CFB (864.12.003), a VENI grant to KVH (016.171.068) and under the Gravitation Programme (024.001.006 supporting AFM). We also gratefully acknowledge funding from the Wellcome Trust UK Strategic Award (098369/Z/12/Z). The funders had no role in study design, data collection and analysis, decision to publish, or preparation of the manuscript.

## Author contributions

AFM, KVH and CFB devised the experiments, AFM and KVH analysed the data and all authors wrote the manuscript.

